# Rewiring PBMC responses to prevent CHIKV infection-specific monocyte subset redistribution and cytokine responses

**DOI:** 10.1101/2020.06.04.132340

**Authors:** José Alberto Aguilar-Briseño, Mariana Ruiz Silva, Jill Moser, Mindaugas Pauzuolis, Jolanda M. Smit, Izabela A. Rodenhuis-Zybert

## Abstract

Infection with the mosquito-borne Chikungunya virus (CHIKV) causes acute or chronic arthritis in humans. Inflammatory responses mediated by monocytes, the primary target cells of CHIKV infection in the blood, are considered to play an important role in CHIKV pathogenesis. A recent study revealed that the acute phase of CHIKV infection is characterized by a monocyte-driven response, with an expansion of the intermediate monocyte (IM) subset. In this study, we adopted a previously established in vitro model of CHIKV infection in peripheral blood mononuclear cells, to elucidate the mechanism and relevance of IM expansion in CHIKV replication and associated inflammatory responses. Our data show that infectious but not replication-incompetent CHIKV increases the frequency of IM and to a lesser extent, non-classical (NM) monocytes while reducing the number of classical monocytes (CM). The increase of IM or NM frequency coincided with the activation of inflammatory response and occurred in the absence of lymphocytes implying that monocyte-derived cues are sufficient to drive this effect. Importantly, priming of monocytes with LPS prevented expansion of IM and NM but had no effect on viral replication. It did however alter CHIKV-induced cytokine signature. Taken together, our data delineate the role of IM in CHIKV infection-specific innate immune responses and provide insight for the development of therapeutic strategies that may focus on rewiring monocyte immune responses to prevent CHIKV-mediated arthralgia and arthritis.

## Introduction

Over the last decade, chikungunya virus (CHIKV) infection has affected millions of people worldwide[1]. The vast majority of infected patients experience a febrile illness with painful joints. A substantial number of these individuals, however, develop persistent arthritis-like manifestations (12-49%)[1]. The exact mechanism underlying CHIKV-mediated pathogenesis is not completely understood, however, clinical and *in vivo* studies describe local joint inflammation, infiltration of monocytes into the synovial cavity and bone resorption due to increased osteoclast activity[1–3]. Monocytes represent important cellular targets of CHIKV replication in the blood[4,5], and tissue infiltrating monocytes and tissue-resident macrophages have been postulated to act as a vehicles and a reservoir for chronic CHIKV RNA/antigens, respectively [6,7].

As innate sentinels of their host, blood monocytes sense invading pathogens and orchestrate innate immune responses to contain their spread as well as setting the scene for the activation of adaptive immune responses[8]. Via a set of pathogen recognition receptors (PRRs), monocytes are able to detect a variety of complex pathogen associated molecular patterns (PAMPS). Engagement of PRRs triggers several cellular signaling cascades resulting in cell differentiation, release of inflammatory mediators and/or programmed cell death[8–10].

Monocytes can be divided into three subsets based on CD14 and CD16 surface expression: classical (CD14++CD16− CM), intermediate (CD14++CD16+, IM) and non-classical (CD14+CD16++, NM)[11–14]. CM are equipped with a set of PRRs and scavenger receptors, recognizing PAMPS thereby removing microorganisms, lipids, and dying cells via phagocytosis. CM also produce a large variety of effector molecules such as cytokines, myeloperoxidase and superoxide, and play a key role in the initiation of inflammation[15]. IM, also known as inflammatory monocytes, selectively traffic to sites of inflammation where they produce inflammatory mediators thereby contributing to local and systemic inflammation[16,17]. IM have the capacity to infiltrate into tissues where they differentiate into inflammatory macrophages and remove PAMPs and cell debris from the microenvironment[18]. NM or patrolling monocytes function as guardians of the vasculature screening for PAMPS. Once activated, they differentiate into anti-inflammatory macrophages to repair damaged tissues[19].

During homeostasis, circulating monocytes have a half-life of about one to three days. CM represent the most prevalent fraction (80-90%) of blood monocytes followed by IM (5%) and NM (2-5%). However, stress caused by infection or tissue damage can lead to changes in monocytes viability, subset redistribution, and their effector functions. For instance, viral infections are usually associated with an increase in the frequency of IM. Yet, depending on the etiologic agent, the frequency of CM and NM varies considerably[15–22]. Indeed, in vitro engagement of innate receptors by for example TLR7/8 agonists has been shown to cause expansion of IM while TLR4 ligand LPS has been shown to further increase the frequency of CM[20]. The pathogen-tailored responses of the monocyte population are thus likely a result of specific combinations of PAMPs and their respective PRRs, however, the underlying mechanisms remain elusive.

Michlmayr and colleagues recently compared the systemic immune signature of CHIKV-infected individuals in an acute and the convalescent stages of the infection[23]. The ex vivo analysis of PBMCs demonstrated that increased IM frequencies are associated with the induction of inflammation in the acute phase CHIKV infection. Moreover, detection of CHIKV E1 antigen primarily in IM suggested that these cells may also contribute to virus replication [23]. The mechanism and the impact of IM increase on viral replication and associated inflammatory responses remain however elusive. Here, we address these using an in vitro model of acute CHIKV infection in primary blood mononuclear cells. By rewiring monocyte responses with different TLR ligands prior to CHIKV exposure, we delineate the role of IM in virus replication and innate responses to CHIKV infection.

## Results

### Active CHIKV infection stimulates expansion of intermediate monocytes

To elucidate the mechanism of monocyte shift during CHIKV infection, we first assessed the contribution of CHIKV particle sensing and/or infection to monocyte subset distribution. We studied monocytes in the context of PBMCs as other cell subsets have previously been shown to be important for the function of monocytes during CHIKV infection[4,5]. PBMCs were isolated from seven donors, as described previously[4] and infected with CHIKV strain LR2006 OPY1 (from hereon called CHIKV) at (multiplicity of infection) MOI 10. The UV-inactivated (UV-CHIKV) was used to evaluate the importance of viral replication in monocytes subset redistribution. At 48 hpi, single cell, live monocytes were analyzed on the basis of CD14 and CD16 expression and gated as CD14 and/or CD16 positive cells using the corresponding isotype controls (Fig 1A). The culture of PBMCs in mock conditions (Fig 1B) altered the baseline distribution of monocyte subsets in freshly isolated PBMC[21] in which CM represented the most abundant subtype followed by IM and NM (S1 Fig). Importantly however, in line with the ex vivo findings[23] CHIKV infection induced a significant, approximately 2-fold expansion of IM and NM frequencies and reduction of CM when compared to the mock condition (Fig 1B and 1C). No significant changes were observed with UV-CHIKV when compared to the mock which suggests that the increase in IM frequency is driven by CHIKV replication. In contrast to CHIKV, exposure of PBMC to LPS (TLR4/CD14 agonist) led to enrichment of CM and almost complete depletion of IM corroborating previous findings[20] (Fig 1D). Furthermore, TLR7/8 agonist R848 and TLR2 agonist PAM3CSK4, also increased the frequency of CM (Fig 1D) rather than that of IM as observed by Kwissa *et al*[20].

**Fig 1.**
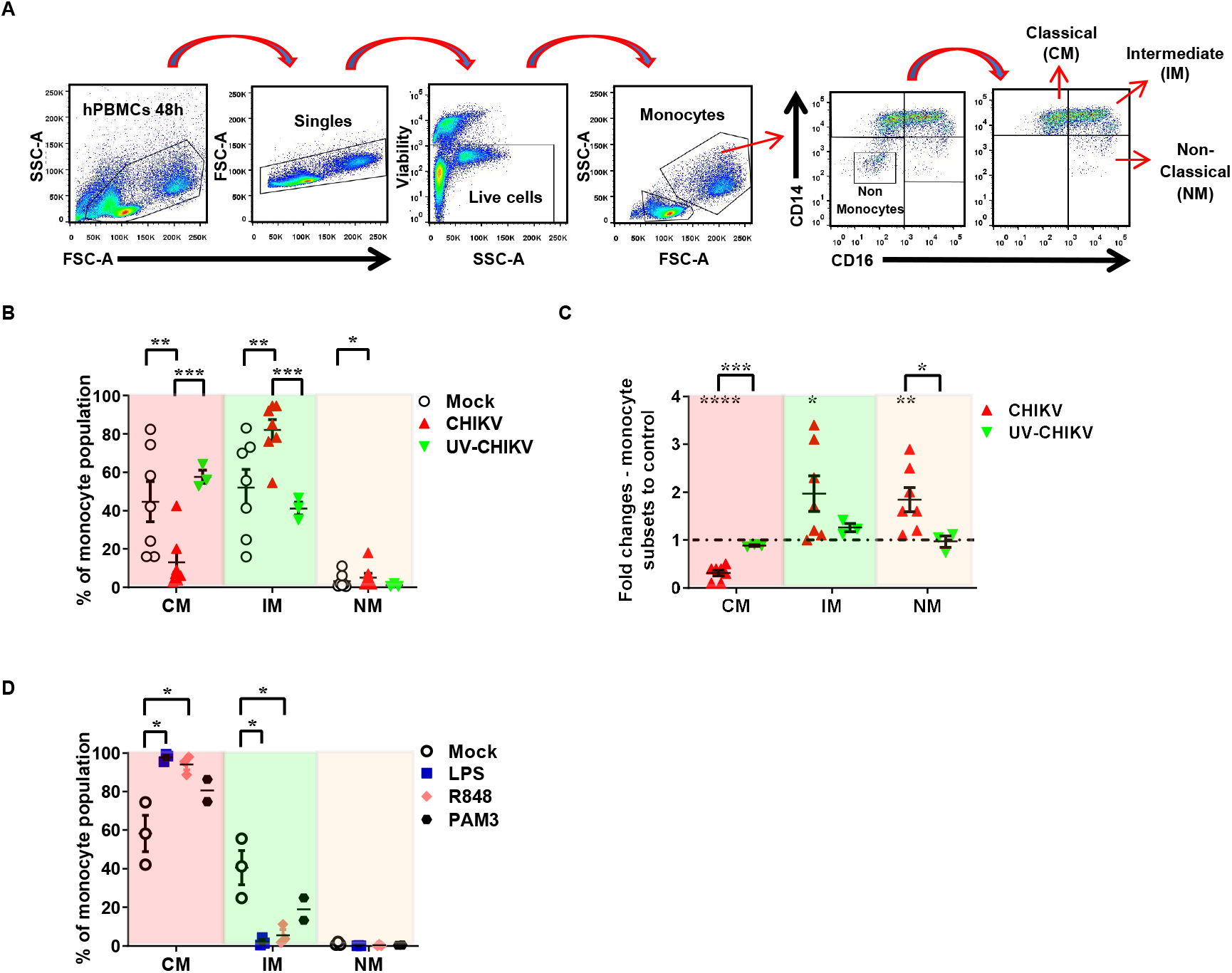
CHIKV infection increases the frequency of intermediate monocytes. (**A-D**) PBMCs from healthy donors (n=3-7) were either infected with CHIKV-LR at MOI of 10 (n=7) or its UV-inactivated equivalent (UV-CHIKV) (n=3), or treated with LPS (100 ng/mL, n=3), R848 (2 μg/mL, n=3), PAM3CSK4 (PAM3, 50ng/mL, n=2) for 48hpi. (**A**) Gating strategy to define monocyte subsets. (**B**) Frequencies of monocyte subsets after infection with CHIKV depicted as percentages or (**C**) as fold changes relative to the mock. (**D**) Frequencies of monocyte subsets after treatment with TLR agonists. CM: classical monocytes, IM: intermediate monocytes and NM: non-classical monocytes. Bar represents mean ± SEM. P values were obtained by paired (**D**) or unpaired (**B** and **C**) one-tailed t test. (*P< 0.05, **P<0.01, ***P<0.001, ****P<0.0001).

Importantly, the increase in IM frequency following CHIKV infection was not the result of a replication-mediated cytopathic effect in PBMCs (S2A, S2B and S2C Fig). In fact, only replication-incompetent UV-CHIKV moderately decreased PBMCs viability (S2A Fig). However, when gated on specific cellular populations, neither CHIVK or UV-CHIKV had an effect on the viability of monocytes and lymphocytes (S2B and S2C Fig). In fact, only LPS showed a considerable cytopathic effect on monocytes but not lymphocytes, as compared to mock (S2B and S2C Fig). Furthermore, the redistribution of monocyte subsets following CHIKV infection was largely independent of cues from other cells since the effect was sustained in PBMCs depleted of T and B lymphocytes, NK and DCs (S3 Fig).

### CHIKV infection-mediated shift in IM coincides with induction of specific inflammatory responses

We next assessed if monocyte redistribution influenced CHIKV-infection specific cytokine signatures (Fig 2A). Initial assessment of gene expression analysis of several inflammatory cytokines from 2 donors showed that replication-competent CHIKV but not the UV-inactivated virus increased the mRNA levels of *IL-6* and *IFN-β* (Fig 2B). On the other hand, UV-CHIKV increased the expression of *IL-8* mRNA by circa 2-fold while replication-competent CHIKV reduced its expression (Fig 2B). Similar mRNA expression signatures were found at 24 hpi for *IL-6, IL-8* and *IFN-β*, but not *TNF-α* (S4 Fig). Subsequent multiplex analysis of cytokines released from infected PBMCs confirmed our gene expression findings for IL-6 and IFN-β (Fig 2C). However, no TNF-α production was detected in the supernatants of infected PBMCs, suggesting that either the observed increase in *TNF-α* mRNA levels did not result in TNF-α production or it is produced later than 48hpi. Unfortunately, IL-8 levels present in all conditions reached the upper limit of quantification, thereby preventing verification of the CHIKV-specific reduction of *IL-8* mRNA. Furthermore, CHIKV but not UV-CHIKV, led to the increase of inflammatory mediators such as IL-10, IP-10 and IFN-α2 (Fig 2C). There was no effect on the gene expression levels of *MAVS* or *IRF3* (S5 Fig), which might suggest that the observed IFN type I production following CHIKV infection was independent of RIG-I/MDA5-mediated sensing of dsRNA[24,25]. We detected no changes in mock levels of TNF-α, IL-1β, IL-12p70, GM-CSF, IFN-γ, IFN-λ1 and IFN-λ2/3 following exposure to (UV-)CHIKV.

**Fig 2.**
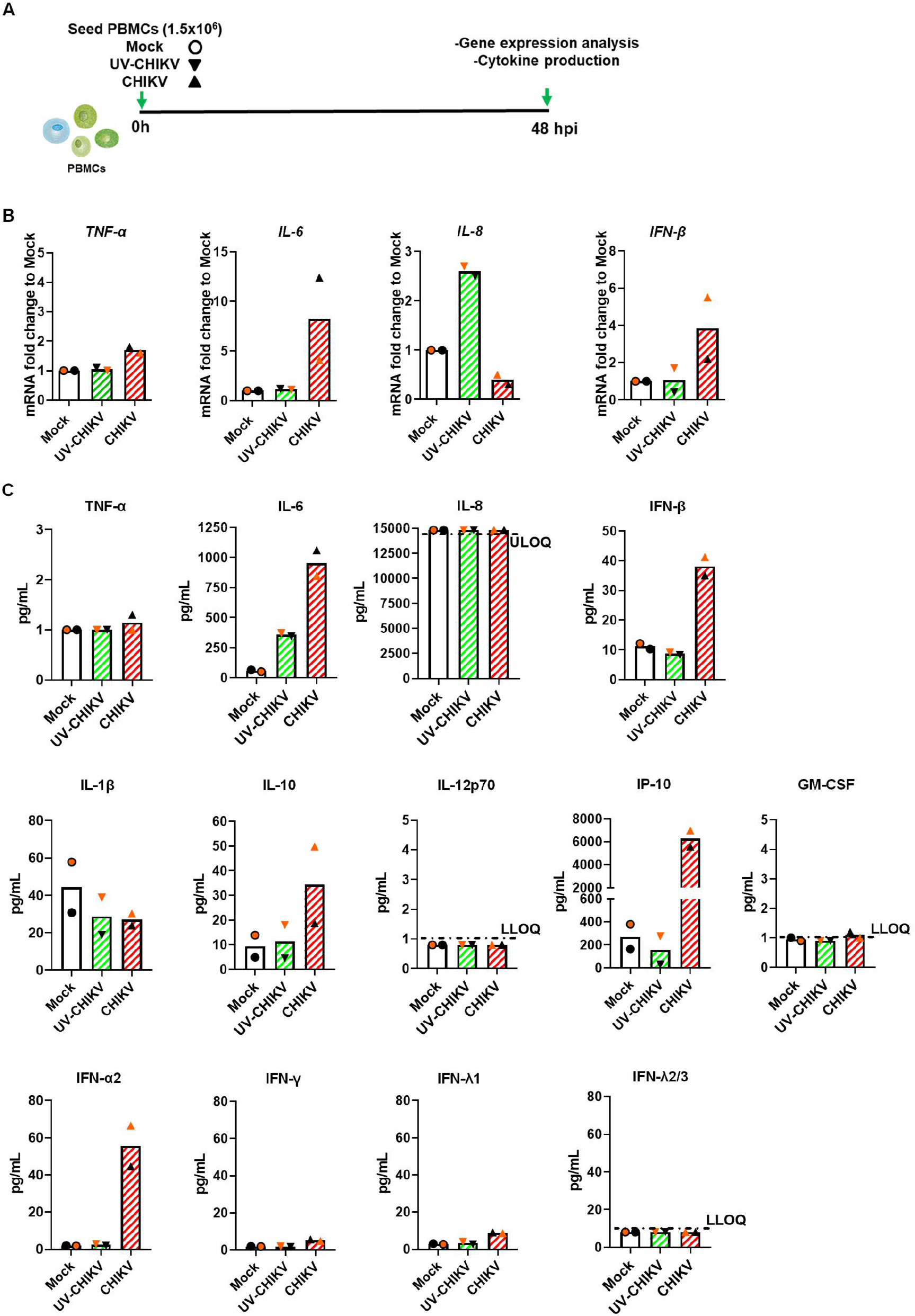
Differential immune responses induced by CHIKV and UV-CHIKV. (**A**) PBMCs from healthy donors (n=2, orange= donor 1, black= donor 2) were (mock)-treated with LPS (600 ng/mL), CHIKV-LR at MOI of 10 or its UV-inactivated equivalent (UV-CHIKV), for 48h. (**B**) Fold changes in gene expression of *TNF-α*, *IL-6*, *IL-8* and *IFN-β* relative to the respective mock. Mock 48h was set as a reference sample. *YWHAZ* was set as a reference gene. (**C**) Concentrations of released cytokines in picograms per milliliter (pg/mL) were determined using LegendPlex. ULOQ and LLOQ (horizontal dotted line) indicate upper and lower limit of quantification, respectively.

Next, we sought to elucidate if CHIKV-mediated responses are specifically associated with an increase in IM and NM frequencies. To this end, we first evaluated the expression levels of the same inflammatory mediators as above but now following the exposure of PBMCs to LPS, which in contrast to CHIKV, increased the frequency of CM (Fig 1D). To avoid a possible bias in expression levels of inflammatory mediators caused by a moderate, albeit consistent, cytopathic effect of LPS on PBMCs at 48hpi (S2 Fig), we evaluated LPS-specific responses at both at 24 hpi and 48hpi (Fig 3 and S6 Fig). Importantly, in line with data from 48hpi, exposure of PBMC to LPS for 24 hpi (hereafter referred to as priming) increased CM frequency and concurrently virtually depleted IM and NM subsets (Fig 3). At the same time, priming had no effect on baseline (mock) mRNA expression levels of *TNF-α* or *IFN-β*, but induced marked 30- and 1000-fold increase in transcription of *IL-8* and *IL-6* genes, respectively (Fig 3 and S6 Fig). Thus, LPS-mediated shift of monocytes subsets and concurrent inflammatory responses were markedly different from those initiated upon CHIKV infection alone.

**Fig 3.**
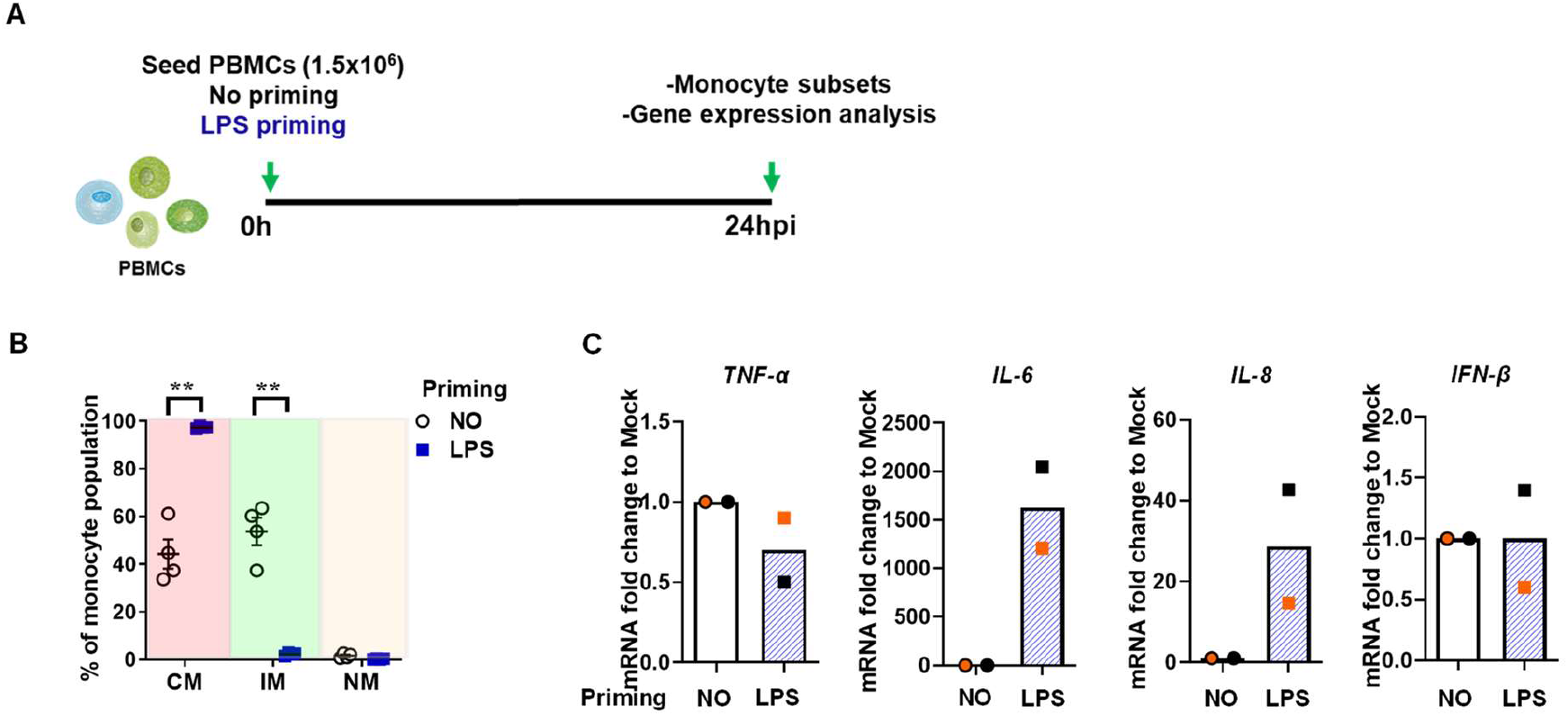
Effect of LPS treatment in cytokines, gene expression and monocyte subsets distribution. (**A**) PBMCs from healthy donors (n=2-4) were (mock)-treated with LPS (600 ng/Ml) for 24h. (**B**) Monocyte subsets distribution as determined by flow cytometry. (**C**) Fold changes in gene expression of *TNF-α, IL-6, IL-8* and *IFN-β* relative to the respective mock. Mock 24h was set as a reference sample. *YWHAZ* was used as a reference gene. Orange symbol=donor 1, black symbol=donor 2, samples from both donors were analyzed in duplicate. Bars represent mean ± SEM. P values were obtained by paired one-tailed t test; **P<0.01.

### Preventing IM expansion alters immune responses but has no effect on CHIKV production

Expansion of a particular monocyte subset often corresponds to the cellular tropism of the virus, as it has been shown for HIV-1, hepatitis C virus (HCV) and Zika virus (ZIKV)[22,26–29]. Since in the acute phase of infection, CHIKV E1 antigen was found primarily in human IM, we first verified that this is also the case in our in vitro system. Indeed, and as expected here considering the prevalence of IM following the infection, CHIKV Ag was present primarily within IM population of PBMCs exposed to replication-competent CHIKV (S7 Fig). Consequently, we next hypothesized that if CHIKV infection leads to the expansion of more susceptible monocyte subsets, priming of the cells with LPS prior to infection would reduce viral replication and likely alter CHIKV-specific immune responses (Fig 4A). Priming PBMCs with LPS prevented subsequent CHIKV infection-mediated increase in IM pool and instead increased CM frequency (Fig 4B). Despite the profound effect on the distribution of monocytes subsets, priming did not influence infectious virus production (Fig 4C). Thus, IM are not more susceptible and/or permissive to CHIKV infection than CM or NM. Rather, due to the substantial increase in the IM population, the likelihood of virus infecting this monocyte population is higher. Priming did however alter the response of PBMC to both UV-CHIKV and infectious virus (Fig 4D). The levels of cytokines including IL-6, IL-8, IFN-β, IL-1β, GM-CSF and IFN-γ were higher in primed and CHIKV-challenged samples when compared to their non-primed counterparts. In contrast, IP-10 and IFN-α2 levels were similar in primed and non-challenged (mock) conditions, indicating that priming overruled and/or prevented CHIKV-specific increases of IP-10 and IFN-α2. Indeed, CHIKV-specific induction of IP-10 and IFN-α2 was reduced in primed conditions. Notably, priming provoked a relatively higher secretion (donor-specific) of IL-6, IL-1β, IFN-β and, too less extent that of GM-CSF and IFN-γ in response to challenge with UV-CHIKV. The priming had no effect on the levels of IFN-λ1 and IFN-λ2 in any condition tested. Altogether, these data suggest that LPS priming alters both monocyte subset distribution and immune responses to CHIKV without affecting virus replication (Fig 5).

**Fig 4.**
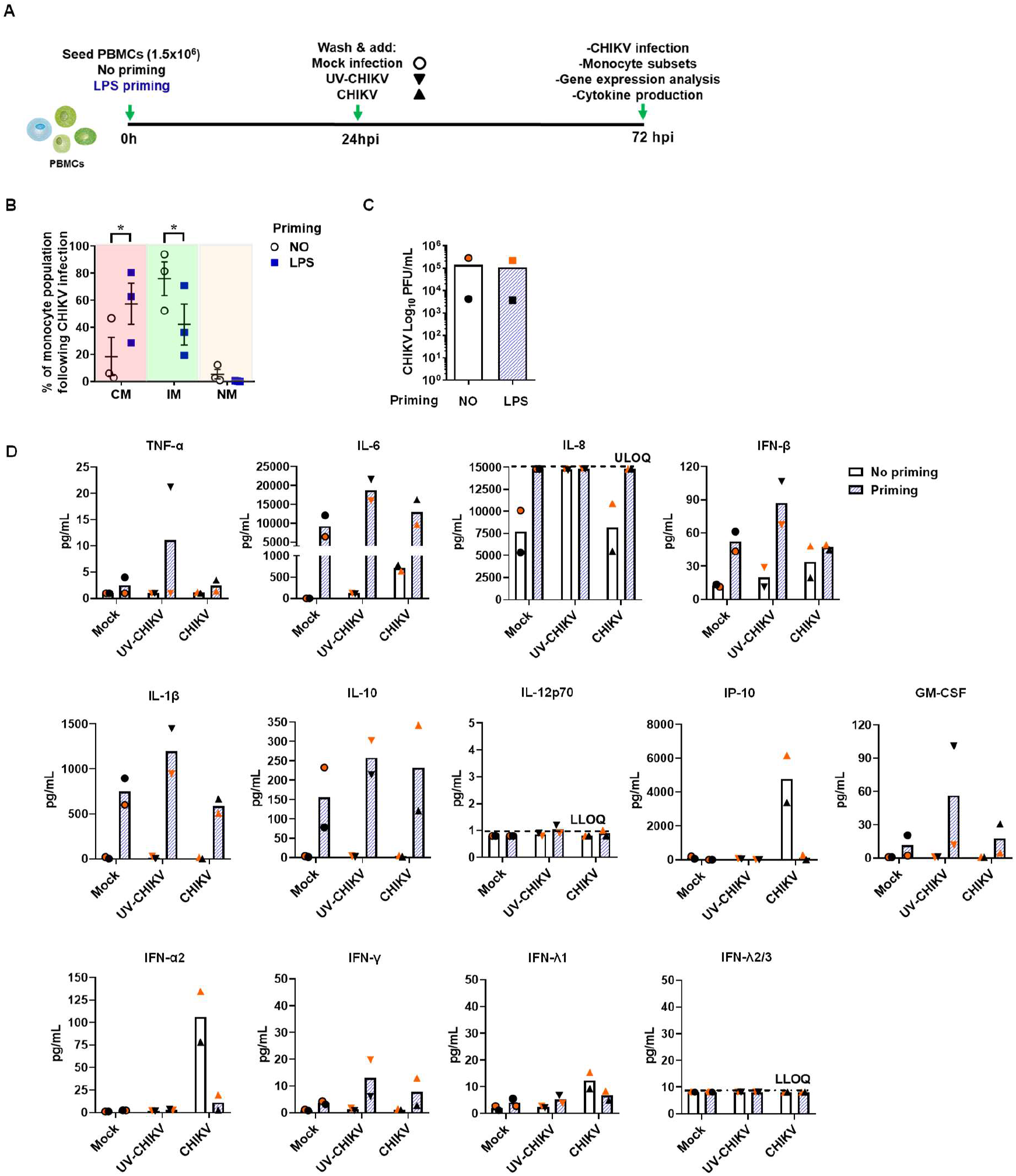
Effect of LPS priming on the infection and innate immune responses following CHIKV infection. (**A**) PBMCs from healthy donors (n=2-3) were (mock)-treated with LPS (600 ng/mL) for 24h and then (mock)-treated with LPS (600 ng/mL), CHIKV-LR at MOI of 10 or its UV-inactivated equivalent (UV-CHIKV). (**B**) Frequencies of monocyte subsets after infection with CHIKV. (**C**) Plaque forming units (PFU) were measured by plaque assay. (**D**) Concentrations of released cytokines in picograms per milliliter (pg/mL) were determined using LegendPlex. ULOQ and LLOQ (horizontal dotted line) indicate upper and lower limit of quantification, respectively. Orange symbol= donor 1, black symbol= donor 2. CM: classical monocytes, IM: intermediate monocytes and NM: non-classical monocytes. Bar represents mean ± SEM. P values were obtained by paired one-tailed t test. (*P< 0.05).

**Fig 5.**
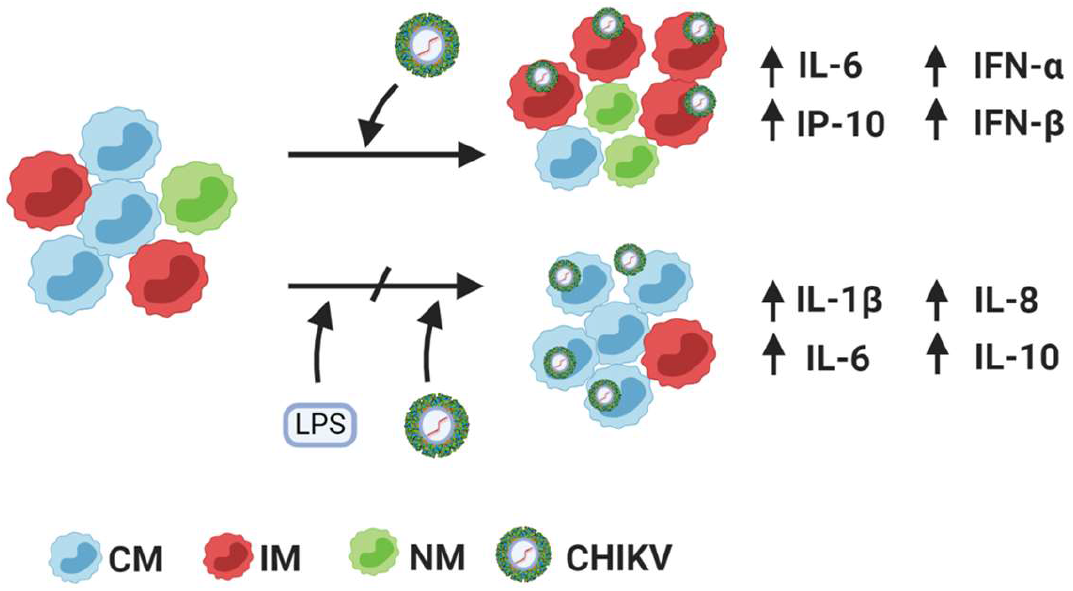
Summary of the results. CHIKV infection of human PBMCs induces an infection-specific innate immune response hallmarked by the increased frequency of intermediate monocytes (IM) and the production of inflammatory mediators such as IL-6, IP-10, IFN-α and INF-β. Priming PBMCs with LPS prevents CHIKV infection-induced monocyte subset redistribution and cytokine responses without altering CHIKV replication. Figure created with BioRender.com

## Discussion

Here, we used a previously established in vitro model of CHIKV infection in peripheral blood mononuclear cells to elucidate the mechanism and effect of IM expansion in the acute phase of CHIKV infection[23] on virus replication and associated inflammatory responses. Our findings indicate that the increase in IM population is dependent on CHIKV replication and occurs irrespectively of cues from other leukocytes. Furthermore, the redistribution of monocytes subsets coincided with the presence of CHIKV Ag in IM and infection-mediated inflammatory responses. By rewiring PBMCs responses with LPS, we could prevent CHIKV infection-specific monocyte subset redistribution and cytokine responses without altering virus replication.

An increase in IM has been also been reported during the course of several other chronic inflammatory diseases such as rheumatoid arthritis (RA)[30–33]. This indicates that the imbalance in the frequency of monocytes contributes to persistent and exacerbated inflammation. Indeed, Chimen *et al.*,[34] suggested that during inflammation, NM and IM transmigrate more rapidly across the endothelium where they produce high levels of TNF-α thereby promoting neutrophil recruitment into the inflamed tissue. In contrast, CM migrate slower than CD16+ monocytes and upon arrival at the site of inflammation decrease neutrophil recruitment by producing IL-6[34]. Consequently, factors preventing expansion of IM by either triggering CM phenotype by LPS priming as presented here, or depleting IM by glucocorticoid therapy, could be applied to decrease inflammation in diseases hallmarked by an expansion of CD16+ monocytes[35]. Importantly, the expansion of IM and NM and concurrent reduction of CM frequency described here in our in vitro model of CHIKV infection, underscores the ability of CM to differentiate into IM and finally to NM as suggested by previous *in vivo* studies[13,36–39]. Importantly, as life spans and migratory function differ between the monocytes subsets[13,17], future studies should elucidate how quickly conversion occurs, and how long each of these subsets remain in circulation following CHIKV infection before infiltrating synovial tissue. Identifying the mechanisms governing the regulation of monocyte differentiation as well as its kinetics and function will be crucial to better understand their role in the pathogenesis of CHIKV-mediated arthritis that will facilitate the development of immunomodulatory therapies.

Expansion of a particular monocyte subset has been shown to correspond to the cellular tropism of many viruses. For instance, HIV-1 causes expansion of IM and NM and these cells are more permissive to the virus than CM, mainly because they express the entry receptors for HIV-1, CCR5 and CD4[22,26]. Likewise, hepatitis C virus (HCV) triggers an increase of IM and NM and preferentially infects these subsets as they express high levels of the virus cell entry receptor CD81[27,28]. Zika virus (ZIKV) causes expansion of the IM and these cells are also the most susceptible monocytes to infection[29]. While we were working on this manuscript, Michlmayr and colleagues reported that CHIKV Ag could be found primarily in IM of infected patients[23]. Based on these observations we hypothesized that altering the numbers of IM would have an effect on virus replication. To investigate this we took advantage of findings which showed opposing effects of TLR4 and TLR2/7/8 ligands[20] on CM and IM frequencies. Importantly, however, our study demonstrates that although CHIKV virus antigen is found in IM, lack of this particular subset and predominance of CM in LPS primed PBMC did not affect virus replication. This indicates that in the case of CHIKV, IM expansion does not represent a pro-viral mechanism, rather an innate response of the host to a particular set of viral PAMPs. Moreover, since the increase in IM population occurred independently of lymphocytes, PRRs expressed on monocytes are likely to govern the mechanisms underlying monocyte differentiation. Finally, given that UV-inactivated CHIKV failed to trigger IM expansion, it is likely that RNA sensing TLRs such as TLR3/7/8 [3,40–42] and/or RLRs[43] play an important role in the process.

Counteracting pro-inflammatory mediators coupled with a simultaneous boost of antiviral responses might be the most successful approach to achieve balanced innate responses that will favour virus containment without causing aberrant pathologies. Accordingly, rewiring inflammatory responses without reducing virus replication, as observed here in LPS-primed PBMCs, is arguably a suboptimal strategy to mitigate CHIKV disease. Interestingly, however, rewiring immune responses in CHIKV-infected mice with bindarit, an anti-inflammatory drug blocking inter alia MCP-1 production[44], reduced joint swelling and prevented bone erosion despite having no effect on viremia levels[45]. Thus, mitigation of disease may be achieved without decreasing viral replication. Indeed, there is no consensus regarding the correlation between peak viremia CHIKV titers and disease progression[23,46]. Since our study did not assess the effect of LPS priming on CHIKV pathogenesis, we cannot, nor do we intend to, equate the effect of LPS found in PBMCs with that found *in vivo* with MCP-1 inhibitor. However, TLR4-mediated reduction of MCP-1 expression in monocytes is among several identified responses in LPS-treated PBMCs[47]. Based on these observations, it would be worthwhile to assess the effect of LPS on CHIKV pathogenesis *in vivo*. Moreover, given that increased levels of LPS in the blood have been found in several inflammatory diseases[48–50], such studies will inform us how co-microbial infections and/or how microbial translocation will influence host responses to CHIKV and ultimately disease progression. In addition, they are imperative to understanding whether pharmacological targeting of inflammatory mechanisms early in infection could serve as a strategy to prevent CHIKV-mediated arthralgia and arthritis, possibly, even without the necessity to decrease viral replication.

In summary, our study provides important insights into the mechanism and role of IM expansion in CHIKV replication and infection-mediated immune responses. Further studies are required to identify receptors involved in monocyte differentiation during CHIKV infection and validate their function in disease progression.

## Methods

### Cells

PBMCs were isolated by standard density gradient centrifugation procedures with Ficoll-Plaque™ Plus (GE Healthcare) from buffy coats obtained with written informed consent from healthy volunteers, in line with the declaration on Helsinki (Sanquin Bloodbank, Groningen, the Netherlands). The PBMCs were cryopreserved at −196 °C. Vero-E6 and Vero-WHO were cultured in DMEM supplemented with FBS, penicillin (100 U/mL), streptomycin (100 μg/mL), HEPES (10 mM) and glutamine (200 mM). All of the cell lines used were tested negative for the presence of Mycoplasma spp. using a commercial functional method (Lonza, the Netherlands) and/or in-house qPCR assay adapted from Baronti *et al.*,[51].

### Monocyte enrichment

Monocytes were isolated from thawed PBMCs using the MagniSort™ human pan-monocyte enrichment kit (eBioscience). Briefly, a single-cell suspension containing 1 × 10^8^ PBMCs per mL of cell separation buffer (PBS, 3% FBS, 10mM EDTA) was prepared. Cells were then incubated for 10 min with 20 μL of MagniSort™ enrichment antibody cocktail per 100 μL of cell suspension. Cells were washed and resuspended in separation buffer. Then 10 μL of MagniSort™ negative selection beads were added per 100 μL of cell suspension. Following 5 min of incubation, a magnet was used to remove the bead-bound non-monocyte lymphocytes from the monocytes. Enrichment efficiency was determined by flow cytometry staining with anti-human CD14 eFluor 450 (clone 613D, eBioscience) and anti-human CD16 APC (clone CB16, eBioscience).

### Virus

CHIKV (La Reunion OPY1) was a gift from A. Merits (University of Tartu, Estonia), and was produced from infection cDNA clones and passaged twice in Vero E6 cells as described before[52]. The infectivity of CHIKV was determined by measuring the number of plaque-forming units (PFU) by standard plaque assay on Vero WHO and the number of genome-equivalent copies (GEc) by quantitative RT-PCR (RT-qPCR), as described previously[53]. Virus inactivation was performed by 1.5h incubation of virus aliquots under UVS-28 8-watt Lamp. Inactivation below level of detection 35 PFU/ml was confirmed using standard plaque assay on Vero-WHO[53]. CHIKV stock used in this study tested negative for Mycoplasma spp. using a commercial functional method (Lonza, the Netherlands) and/or in-house qPCR assay adapted from Baronti *et al.*,[51].

### In vitro infections

PBMCs (1×10^6^ cells/mL) were treated with PAM3CSK4 (50 ng/mL, InvivoGen), LPS (*E.coli* K12, 100 ng/mL, InvivoGen), R848 (2 μg/mL, InvivoGen), CHIKV-LR at a multiplicity of infection (MOI) of 10 or its UV-inactivated equivalent (UV-CHIKV-LR) for 48h. Monocytes (2×10^5^ cells) were infected with CHIKV-LR at MOI 10 or its equivalent UV-CHIKV-LR for 48h. To evaluate the effect of the LPS priming on CHIKV infection, PBMCs were (mock)-primed with LPS for 24h and then (mock)-treated with LPS or infected with CHIKV-LR at MOI 10 for 48h or its equivalent UV-CHIKV-LR for 48h. For viral production, CHIKV-LR infected cells were washed after infection and media replenished (600 μL), 48h later cell-free supernatants were collected and preserved at −80°C. For gene expression and cytokine production analysis cell-free supernatants and cell lysates were collected at 48h, 24h priming and 48h post-priming, samples were preserved at −80°C and –20°C, respectively.

### Flow cytometry analysis

Surface expression of CD14 and CD16 was measured on PBMCs and enriched monocytes directly after thawing of the cells, after infection with CHIKV or treatment with the agonists but prior to infection and 24h and 48h post infection/treatment with the agonists. Briefly, PBMCs were stained with Fixable Viability Dye eFluor 780, anti-human CD14 eFluor 450 (clone 613D) and anti-human CD16 APC (clone CB16). All purchased from eBioscience. Samples were measured on a FACSverse flow cytometer (BD Biosciences). Isotype-matched antibodies labelled eFluor 450 (clone P3.6.2.8.1) and APC (clone P3.6.2.8.1), both eBioscience, were used as controls to compare the expression of each marker. To measure the number of infected monocytes, PBMCs were fixed at 48hpi and stained intracellularly using CHIKV E1-specific rabbit antibody (kindly gifted by dr. G. Pijlman from Wageningen University) and secondary chicken anti-rabbit AF647 (Life Technologies). Samples were measured on Aurora flow cytometer (Cytek®). Data were analyzed using the Flowjo software (BD Biosciences).

### Gene expression analysis

Total RNA was isolated from PBMCs using a RNeasy Plus Mini Kit (Qiagen, Leusden, The Netherlands), according to the manufacturer’s instructions. RNA integrity was checked, cDNA synthesized, and RT-qPCR performed using the ViiA 7 system (Applied Biosystems/ThermoFisher Scientific) as previously described[54]. Assay on demand primers were from Applied Biosystems (Nieuwerkerk aan de IJssel, The Netherlands) included *IL-6* (Interleukin-6, Hs00174131_m1), *IL-8* (Interleukin-8, Hs00174103_m1), *TNF-α* (Tumor necrosis factor alpha, Hs00174128_m1), *IFN-β* (Interferon beta, Hs01077958_s1), *IRF3* (IFN regulatory factor 3, Hs01547283_m1) and *MAVS* (Mitochondrial antiviral signaling protein, Hs00920075_m1). Duplicate real-time PCR analyses were performed for each sample, and the obtained threshold cycle (CT) values were averaged. Gene expression was normalized to the expression of housekeeping gene (*YWHAZ*, Hs01122445_g1) resulting in the ΔCT value. The relative mRNA level was calculated by 2^− ΔCT^.

### Multiplex analysis of cytokines and chemokines

Human anti-virus response panel (13-plex, LEGENDplex™, BioLegend) was used to determine the protein levels of IL-1β, TNF-α, IL-6, IL-8, IL-10, IL-12p70, IP-10, GM-CSF, IFN-α2, IFN-β, IFN-γ, IFN-λ1 and IFN-λ2/3. Data were collected using a FACSverse flow cytometer (BD Biosciences) and analyzed with LEGENDplex™ v8.0 (BioLegend).

### Statistical analysis

Data analysis was performed using Prism 6.01 (Graphpad, USA). Data are shown as mean ± SEM. Paired one-tailed t-test or unpaired one-tailed t-test were used to determine statistical significance. In all tests, values of *p<0.05, **p<0.01, ***p<0.001 and ****p<0.0001 were considered significant.

## Supporting information

**S1 Fig. Changes in the monocyte subsets distribution due to in vitro culturing.** PBMCs from healthy donors (n=1) were cultured for 48h. Frequencies of monocyte subsets were determined by flow cytometry immediately after thawing and at 48h of culture.

**S2 Fig. Cell viability at 48h post-treatment.** PBMCs from healthy donors (n=2-4) were (mock)-treated with LPS (100 ng/mL, n=2), CHIKV-LR at MOI of 10 (n=4) or its UV-inactivated equivalent (UV-CHIKV) (n=4), for 48h. PBMCs were collected and stained with fixable viability dye. (**A**) Viability of total PBMCs, (**B**) monocytes and (**C**) lymphocytes. Bar represents mean ± SEM. P values were obtained by unpaired one-tailed t test (*P<0.05, ** P<0.01).

**S3 Fig. Active CHIKV infection increases intermediate monocytes’ frequency.** Monocytes were enriched by negative selection from PBMCs isolated from healthy donors (n=3) and then (mock)-infected with CHIKV-LR at MOI of 10 for 48h. (**A**) Enrichment efficiency as assessed by flow cytometry (**B**) Frequencies of monocyte subsets were determined by flow cytometry after infection. CM: classical monocytes, IM: intermediate monocytes and NM: non-classical monocytes. Bar represents mean ± SEM. P values were obtained by paired one-tailed t test (*P< 0.05).

**S4 Fig. Differential expression of cytokines after 24h of infection with CHIKV.** PBMCs from healthy donors (n=2, orange= donor 1, black= donor 2) were (mock)-infected with CHIKV-LR at MOI of 10 for 24h. Fold changes in gene expression of *TNF-α, IL-6, IL-8* and *IFN-β* relative to the respective mock. Mock 24h was set as a reference sample. *YWHAZ* was used as a reference gene.

**S5 Fig. Gene expression levels of *IRF3* and *MAVS* after 24h and 48h of culture.** PBMCs from healthy donors (n=2, orange= donor 1, black= donor 2) were cultured for 24h and 48h. Fold changes in gene expression of *IRF3* and *MAVS* relative to the respective mock. 0h was set as a reference sample. *YWHAZ* was used as a reference gene.

**S6 Fig. Differential expression of cytokines induced by LPS.** PBMCs from healthy donors (n=2) were (mock)-treated with LPS (100 ng/mL, n=2) for 48h. Fold changes in gene expression of *TNF-α, IL-6, IL-8* and *IFN-β* relative to the respective. Mock 48h was set as a reference sample. YWHAZ was used as a reference gene.

**S7 Fig. CHIKV replicates primarily in IM.** PBMCs from healthy donors (n=1) were (mock)-infected with CHIKV-LR at MOI of 20 or its UV-inactivated equivalent (UV-CHIKV) for 48h. Frequencies of CHIKV positive cells and monocyte subsets distribution were determined by flow cytometry.

## Acknowledgments

We thank Heidi Ende-Metselaar for her excellent technical assistance. We thank Peter Zwiers (Endothelial Biomedicine & Vascular Drug Targeting group, UMCG) for his assistance and guidance in the gene expression experiments. We are grateful to Geert Mesander (Flow Cytometry Unit, UMCG) for his technical expertise and guidance in the flow cytometry assays.

## Author contributions

**Conceptualization:** José Alberto Aguilar Briseño, Mariana Ruiz Silva, Izabela A Rodenhuis-Zybert

**Data curation:** José Alberto Aguilar Briseño, Mariana Ruiz Silva

**Formal analysis:** José Alberto Aguilar Briseño, Mariana Ruiz Silva, Izabela A Rodenhuis-Zybert.

**Funding acquisition:** José Alberto Aguilar Briseño, Mariana Ruiz Silva, Izabela A Rodenhuis-Zybert.

**Investigation:** José Alberto Aguilar Briseño, Mariana Ruiz Silva, Mindaugas Pauzuolis.

**Methodology:** José Alberto Aguilar Briseño, Izabela A Rodenhuis-Zybert.

**Project administration:** Izabela A Rodenhuis-Zybert.

**Resources:** Jill Moser, Jolanda M Smit, Izabela A Rodenhuis-Zybert.

**Supervision:** Izabela A Rodenhuis-Zybert.

**Validation:** José Alberto Aguilar Briseño, Mariana Ruiz Silva, Izabela A Rodenhuis-Zybert.

**Visualization:** José Alberto Aguilar Briseño.

**Writing – original draft:** José Alberto Aguilar Briseño, Mariana Ruiz Silva, Izabela A Rodenhuis-Zybert.

**Writing – review & editing:** José Alberto Aguilar Briseño, Jill Moser, Jolanda M Smit, Izabela A Rodenhuis-Zybert.

## Data Availability Statement

All relevant data are within the manuscript and its supplementary information files.

## Funding

JAAB was supported by de Cock – Hadders Stichting grant and by CONACyT, Mexico. MRS was supported by de Cock – Hadders Stichting grant. IARZ was supported by Research Grant 2019 from the European Society of Clinical Microbiology and Infectious Diseases (ESCMID). Funding agencies had no role in the experimental design, decision to publish, or preparation of the manuscript.

## Competing interests

The authors declare that they have no competing interests.

## Notes

### Competing Interest Statement

The authors have declared no competing interest.

