## Supporting information for "Rewiring PBMC responses to prevent CHIKV infection-specific monocyte subset redistribution and cytokine responses"

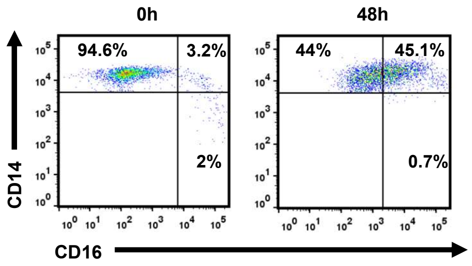


**S1 Fig. Changes in the monocyte subsets distribution due to in vitro culturing.** PBMCs from healthy donors (n=1) were cultured for 48h. Frequencies of monocyte subsets were determined by flow cytometry immediately after thawing and at 48h of culture.

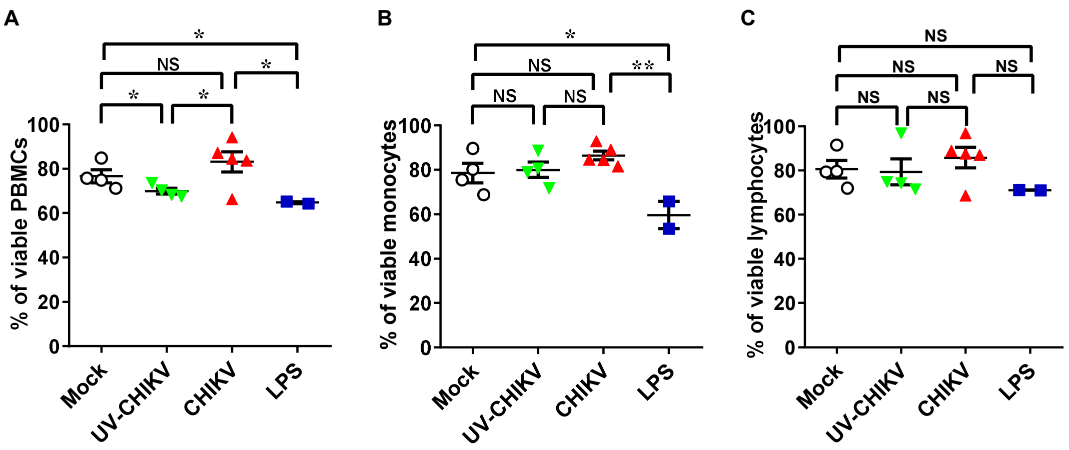


**S2 Fig. Cell viability at 48h post-treatment.** PBMCs from healthy donors (n=2-4) were (mock)-treated with LPS (100 ng/mL, n=2), CHIKV-LR at MOI of 10 (n=4) or its UV- inactivated equivalent (UV-CHIKV) (n=4), for 48h. PBMCs were collected and stained with fixable viability dye. (**A**) Viability of total PBMCs, (**B**) monocytes and (**C**) lymphocytes. Bar represents mean ± SEM. P values were obtained by unpaired one-tailed t test (*P<0.05, ** P<0.01).

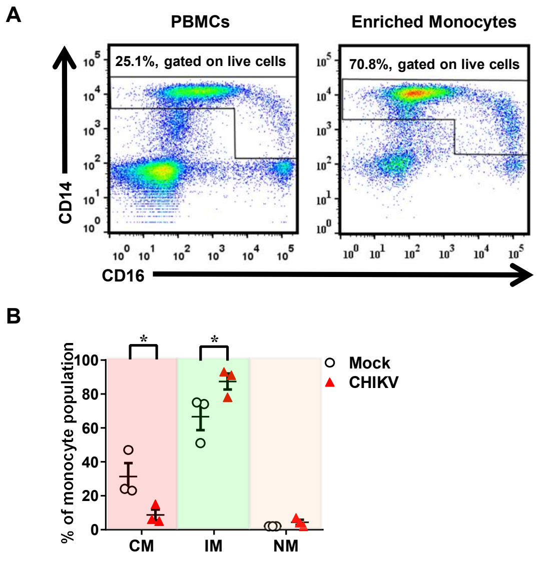


**S3 Fig. Active CHIKV infection increases intermediate monocytes' frequency.** Monocytes were enriched by negative selection from PBMCs isolated from healthy donors (n=3) and then (mock)- infected with CHIKV-LR at MOI of 10 for 48h. (**A**) Enrichment efficiency as assessed by flow cytometry (**B**) Frequencies of monocyte subsets were determined by flow cytometry after infection. CM: classical monocytes, IM: intermediate monocytes and NM: non-classical monocytes. Bar represents mean ± SEM. P values were obtained by paired one-tailed t test (*P< 0.05).

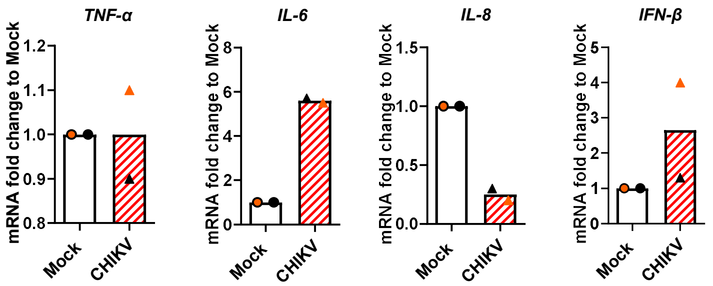


**S4 Fig. Differential expression of cytokines after 24h of infection with CHIKV.** PBMCs from healthy donors (n=2, orange= donor 1, black= donor 2) were (mock)-infected with CHIKV-LR at MOI of 10 for 24h. Fold changes in gene expression of *TNF-α*, *IL-6*, *IL-8* and *IFN-β* relative to the respective mock. Mock 24h was set as a reference sample. *YWHAZ* was used as a reference gene.

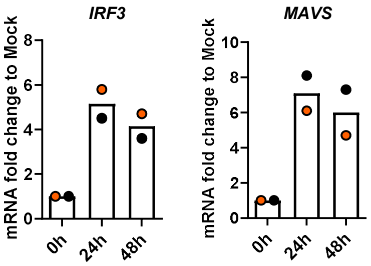


**S5 Fig. Gene expression levels of *IRF3* and *MAVS* after 24h and 48h of culture.** PBMCs from healthy donors (n=2, orange= donor 1, black= donor 2) were cultured for 24h and 48h. Fold changes in gene expression of *IRF3* and *MAVS* relative to the respective mock. 0h was set as a reference sample. *YWHAZ* was used as a reference gene.

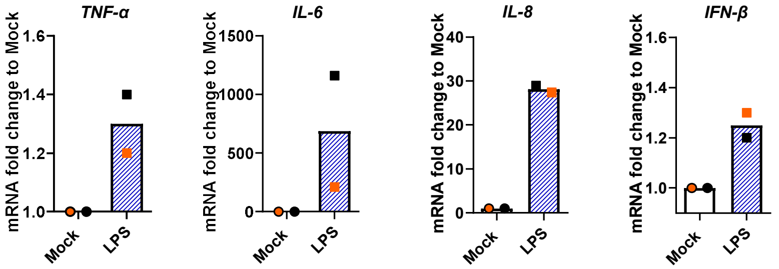


**S6 Fig. Differential expression of cytokines induced by LPS.** PBMCs from healthy donors (n=2) were (mock)-treated with LPS (100 ng/mL, n=2) for 48h. Fold changes in gene expression of *TNF-α*, *IL-6*, *IL-8* and *IFN-β* relative to the respective. Mock 48h was set as a reference sample. *YWHAZ* was used as a reference gene.

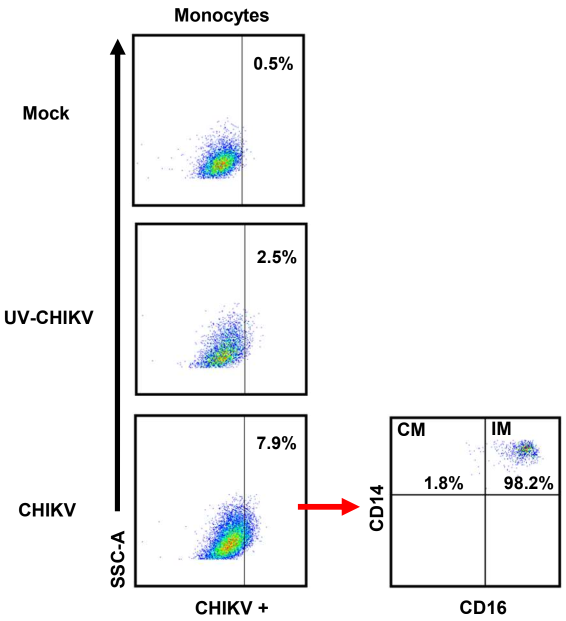


**S7 Fig. CHIKV replicates primarily in IM.** PBMCs from healthy donors (n=1) were (mock)-infected with CHIKV-LR at MOI of 20 or its UV- inactivated equivalent (UV-CHIKV) for 48h . Frequencies of CHIKV positive cells and monocyte subsets distribution were determined by flow cytometry.
